# Fission yeast Smi1p participates in the synthesis of the primary septum by regulating β-1,3-glucan synthase Bgs1p function

**DOI:** 10.1101/2022.05.23.493069

**Authors:** Kriti Sethi, Juan C. G. Cortés, Mamiko Sato, Masako Osumi, Naweed I. Naqvi, Juan Carlos Ribas, Mohan Balasubramanian

**Affiliations:** Mechanobiology Institute, National University of Singapore, 5A Engineering Drive 1, Singapore 117411; Temasek Life Sciences Laboratory, National University of Singapore, 1 Research Link, Singapore, 117604; Instituto de Biología Funcional y Genómica, Consejo Superior de Investigaciones Científicas (CSIC) and Universidad de Salamanca, Salamanca, Spain; Laboratory of Electron Microscopy, Japan Women’s University, Mejirodai, Bunkyo-ku, 102-8681, Tokyo, Japan; Department of Chemical and Biological Sciences, Faculty of Sciences, Japan Women’s University, Mejirodai, Bunkyo-ku, 102-8681, Tokyo, Japan; NPO Integrated Imaging Research Support, Hirakawa-cho, Chiyoda-ku, Tokyo 102-0093, Japan; Japan Women’s University, Mejirodai, Bunkyo-ku, 102-8681, Tokyo, Japan; Department of Biological Sciences, National University of Singapore, Singapore 117543; Centre for Mechanochemical Cell Biology, Division of Biomedical Sciences, Warwick Medical School, University of Warwick, Coventry, UK, CV4 7AL

**Keywords:** cell wall synthesis, division septum, glucan synthesis, cytokinesis

## Abstract

Cytokinesis is the concluding step of the cell cycle. Coordination between multiple cellular processes is essential for the success of cytokinesis. The fission yeast, *Schizosaccharomyces pombe*, like other fungal cells is contained within a cell wall. During cell division, the external cell wall is extended inwards to form a special septum wall structure in continuity with the cell wall. The primary septum, the central component of the three-layered division septum, is enriched with linear β-1,3-glucan formed by Bgs1p, a β-1,3-glucan synthase. In this study we uncover a novel essential protein, Smi1p, that functions as a suppressor of the Bgs1p temperature-sensitive mutant, *cps1-191*. We observe a rescue in the cell wall composition and ultrastructure and also in actomyosin ring dynamics. Further, we identify a colocalization and physical association between Bgs1p and Smi1p. Altogether, our results indicate that Smi1p regulates the function of Bgs1p during cytokinesis.

## 1. Introduction

The fungal cell wall serves as an essential barrier to maintain cell shape and integrity, and remodelling of the cell wall through the cell cycle is essential for viability (Cabib & Arroyo, 2013; Lesage et al., 2005). In *Schizosaccharomyces pombe*, during cytokinesis, a special three layered cell wall containing division septum is deposited centripetally in coordination with actomyosin ring constriction (Cortés et al., 2016; Sipiczki, 2007). Whereas the actomyosin ring positions the septum as well as generates tension for ingression of the plasma membrane, new membranes and the septum physically divide the mother cell into two daughters (Ramos et al., 2019). The division septum contains a primary septum flanked by secondary septa on both sides. The protein Bgs1p/Cps1p synthesizes the linear β-1,3-glucan at the primary septum (Cortés et al., 2002, 2007; Ishiguro et al., 1997; Le Goff et al., 1999; Liu et al., 1999, 2000). The glucan synthase Bgs1p acts in concert with other proteins including other α-1,3-glucan and β-1,3-glucan synthases to ensure synthesis of a well-defined three layered division septum (Cortés et al., 2012, 2015). Of these, Bgs4, encodes another β-1,3-glucan synthase synthesizing branched β-1,3-glucan, while Ags1/Mok1, encodes α-1,3-glucan synthase synthesizing α-1,3-glucan (Cortés et al., 2012; Muñoz et al., 2013). As such, both, Bgs4 and Ags1, play an important role in secondary septum assembly as well as in primary septum and cell integrity. Other proteins, such as F-BAR protein Cdc15p and the secretory apparatus have been implicated in the transport of Bgs1p to the division site at the time of septation (Arasada & Pollard, 2014; Liu et al., 2002).

The budding yeast Knr4p/Smi1p was identified in a screen for resistance to *Hansenula mrakii* killer toxin K9 (Hong et al., 1994). Knr4p was also isolated as a suppressor for cell wall mutants, which are hypersensitive to calcofluor white, with Knr4p’s phosphorylation being essential for this rescue (Basmaji et al., 2006; Ficarro et al., 2002; Martin et al., 1999). Upon overexpression, Knr4p represses expression of the chitin synthase genes (Martin et al., 1999). Though Knr4p is non-essential in budding yeast, the null mutant displays a delay in growth at small budded stage and later shows an abnormal bud neck (Ohtani et al., 2004; Ohya et al., 2005). Further, the Knr4p null mutant exhibits synthetic lethality with other morphogenesis and cell wall biogenesis mutants indicating the important role it plays in cell morphogenesis and division (Costanzo et al., 2010; Lesage et al., 2005). Knr4p is an intrinsically disordered protein, a trait that is believed to allow for its numerous physical interactions (Basmaji et al., 2006; Durand et al., 2008; Martin-Yken et al., 2016). Knr4p functions at the junction between two major signalling pathways in *Saccharomyces cerevisiae* – the cell wall integrity pathway (CWI) and the calcium-calcineurin pathway, either as a coordinator between these two pathways or as a common scaffolding protein (Dagkessamanskaia et al., 2010; Martin-Yken et al., 2003). Further, Knr4p has a very high number of known interacting partners - known physical interaction with 38 different proteins and known genetic interactions with 275 proteins. The identified interactors are involved in various pathways but are all related to morphogenesis and stress response in budding yeast. Knr4p localizes as foci at the bud site, both presumptive and upon emergence (Martin et al., 1999). These foci then relocalize to the division site at the mother-daughter bud neck. In *Candida albicans*, Smi1p contributes to the production of glucan in the biofilm that offers resistance to drugs by sequestration (Nett et al., 2011).

In this study, we identified *S. pomb*e Smi1p, related in sequence to its budding yeast counterpart, to be a suppressor of the Bgs1p temperature-sensitive mutant, *cps1-191*. Our analysis suggests that the two proteins physically interact with each other and that Smi1p might be directly involved in β-1,3-glucan synthesis or might be involved in the transport of Bgs1p to the division site. Our work, therefore identifies a new direct role for Smi1p in primary septum assembly.

## 2. Materials and Methods

### Yeast Strains and methodology

Table S1-S3 lists out *S. pombe* strains, plasmids and primers used in this study. Standard genetic and molecular biology protocols for *S. pombe* were used as described previously (Moreno et al., 1991). For the generation of Smi1p-GFP, primers (MOH5904 and MOH5905) were used to generate GFP-KanMX6 fragments (template pCDL1082) with overhangs to allow for integration at the C-terminus of Smi1p upon transformation into wild type cells. Colonies were selected for geneticin G418 resistance, checked for fluorescence and confirmed by sequencing.

### Plasmids and Recombinant DNA Methods

For the experiment described in Figure 3C and 3D, the plasmid pCDL1000 was modified to express histidine and the leucine expressing fragment was removed. This plasmid is referred to as pEmpty_his3 (pBac6). Next, s*mi1*^*+*^ ORF with 1kb of 5’UTR and 3’UTR was inserted into pBac6 to generate pSmi1_his3 (pBac8) plasmid.

### Microscopy

For the experiment described in Figure 1D, calcofluor white at a final concentration of 10μg/ml (stock:10mg/mL in water) and FITC-ConA at a final concentration of 20μg/ml (stock: 1mg/mL in PBS) were used as described previously (Cortés et al., 2012). A fluorescence microscope (model DM RXA; Leica), a PL APO 63×/1.32 oil PH3 objective, a digital camera (model DFC350FX; Leica) and CW4000 cytoFISH software (Leica) were used to acquire the images and fluorescence intensity was quantified as described (Cortés et al., 2015).

**Figure 1:**
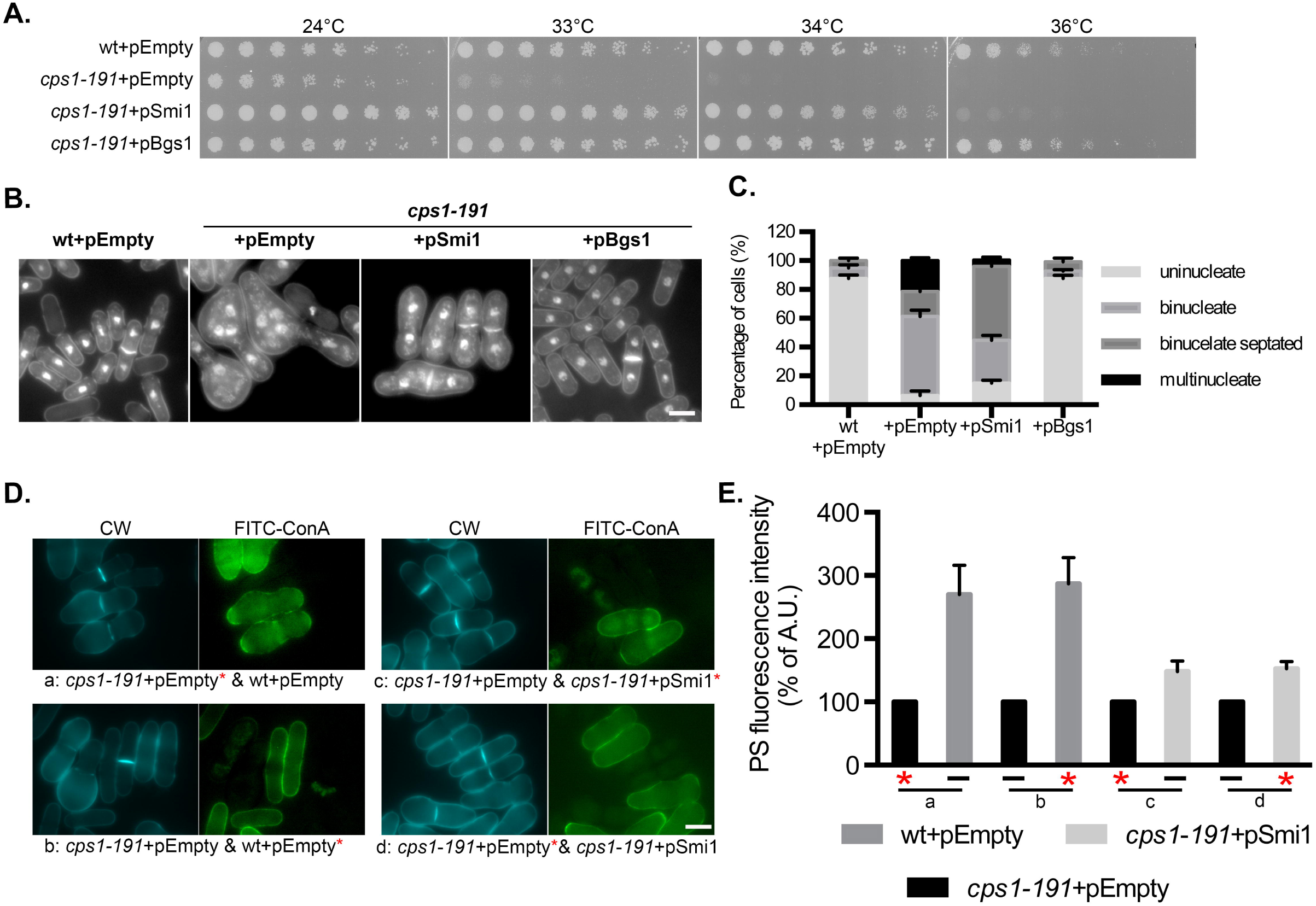
Multi-copy expression of *smi1*^*+*^ rescues glucan synthase *cps1-191* mutant. (A) Spot assay comparing the viability of the strains of indicated genotypes. Cultures of the strains: wt+pEmpty (MBY8558), *cps1-191+*pEmpty (MBY8944), *cps1-191+*pSmi1 (MBY8945) and *cps1-191+*pBgs1 (MBY8947) were grown overnight at 24°C, serially diluted in two-fold steps, spotted on minimal media agar plates without leucine and incubated at various growth temperatures. (B) Aniline blue and DAPI images of medial plane of cells of indicated genotype wt+pEmpty (MBY8558), *cps1-191+*pEmpty (MBY8944), *cps1-191+*pSmi1 (MBY8944) and *cps1-191+*pBgs1 (MBY8947), fixed after shift to 34°C for 16 hr. Scale bar 5μm. (C) Quantification of phenotypes observed in (A). *cps1-191*+pEmpty vs *cps1-191+p*Smi1 binucleate septated cells: p-value=0.0006 (two-tailed T-test) (n=3,= ≥200cells). (D) Calcofluor white (CW) images of the indicated strains: wt+pEmpty (MBY8558), *cps1-191+*pEmpty (MBY8944) and *cps1-191+*pSmi1 (MBY8945) after 16 hr at 34°C, with one strain also stained with concanavalin A (ConA) conjugated with FITC (labelled strain indicated with an asterisk*). a-d refer to the different combinations analyzed and are stated in the figure. (E) Quantification of observed CW fluorescence intensity for experiment done in (B). Percentage of primary septum (PS) fluorescence intensity normalized to septum length of each strain compared to that of *cps1-191*+pEmpty strain, which was assigned a value of 100%. (n≥34cells). Scale bar 5μm. Error bars indicate S.D.

Bright-field images were acquired with an Olympus microscope (IX71; Plan Apochromat 100×/1.45 NA oil objective lens) equipped with a charge-coupled device camera (CoolSNAP HQ; Photometrics) and MetaMorph (v6.2r6) software (Molecular Devices). Spinning-disk images were obtained with a *micro*LAMBDA spinning disk using a microscope (Eclipse Ti; Nikon; Plan Apochromat VC 100×/1.40 NA oil objective lens) equipped with a spinning-disk system (CSUX1FW; Yokogawa Corporation of America), camera (CoolSNAP HQ^2^), and MetaMorph (v7.7.7.0) software. A 491-nm diode-pumped solid-state (DPSS) laser (Calypso), 515-nm DPSS laser (Fandango; Cobolt), and 561-nm DPSS laser (Jive) were used for excitation. Spinning disk confocal images were acquired at 0.5μm step size, for a range of 6μm. All images are presented as 2D maximum intensity projection. Image processing was done using ImageJ (v1.47).

### Transmission electron microscopy

Cells were prepared for TEM as described in (Cortés et al., 2015; Konomi et al., 2003; Osumi & Sando, 1969). Briefly, cells were fixed with 2% glutaraldehyde EM (GA; Electron Microscopy Science) in 50mM phosphate buffer pH 7.2, 150mM NaCl (PBS) for 2h at 4°C, post-fixed with 1.2% potassium permanganate overnight at 4°C. Cells were then embedded in 2% low-melting-point agarose, dehydrated through an ethanol series, and passed through QY-2 (methyl glycidyl ether; Nisshin EM, Tokyo, Japan). Next, cells were embedded in Quetol 812 mixture (Nisshin EM Tokyo, Japan). Ultrathin sections were stained in 4% uranyl acetate and 0.4% lead citrate, and viewed with a TEM JEM-1400 (JEOL, Tokyo, Japan) at 100 kV.

### Multi-copy suppressor screen of *cps1-191*

Mutant *cps1-191* cells were transformed with pTN-L1, a *S. pombe* genomic library (Nakamura et al., 2001), and selected for growth on minimal media plates lacking leucine at 24°C. Viable colonies at 34°C were selected after replicating to YES plates with Phloxin B. The suppressors for *cps1-191* lethality were identified from plasmids in viable colonies. The rescuing gene fragment was identified by sequencing the plasmids (primers MOH1207 and MOH1208).

### Radioactive labelling and fractionation of cell wall polysaccharides

Cell wall analysis was performed as described previously (Ishiguro, 1998; Pérez & Ribas, 2004). Exponentially growing cells at 25ºC in minimal medium lacking leucine were diluted and supplemented with D-[^14^C]-glucose (3μCi/ml), maintained at 24ºC for 24 hr or shifted to 34ºC for 16 hr (Figure S1 and S2). Harvested cells were supplemented with unlabeled cells as carrier, washed twice with 1mM EDTA, and resuspended in 1mM EDTA. Two aliquots of cells were added to liquid scintillation cocktail and total D-[^14^C]-glucose incorporation in the cells was assessed from this. Cell walls were purified from bead-beaten lysed cells by repeated washing and differential centrifugation (once with 1mM EDTA, twice with 5M NaCl, and three times with 1mM EDTA) at 1,500*g* for 5 min. Purified cell walls were heated at 95°C for 30 min and D-[^14^C]-glucose incorporation in the cell walls was monitored in two aliquots. One part of cell wall samples was extracted with 6% NaOH for 60 min at 80°C. The galactomannan fraction was precipitated from the supernatant with Fehling’s reagent by adding unlabeled yeast mannan (4mg) as the carrier. Three volumes of Fehling’s reagent were added to the sample and allowed to precipitate galactomannan overnight at 4°C. Pellets were obtained by centrifugation at 4,000g for 10 min, washed with Fehling’s reagent and solubilized in few drops of 6N HCl. The galactomannan fraction was determined from this after addition of 50mM Tris-HCl, pH 7.5, washing again with the same buffer and mixing both washes, and measuring the radioactivity in a scintillation counter (Perkin Elmer). Second part of cell wall suspensions was incubated with Zymolyase 100T (AMS Biotechnology; MP Biomedicals) in 50mM citrate-phosphate buffer (pH 5.6) for 24h at 37°C, using untreated samples as control. Pellets from centrifugation were resuspended in 1mM EDTA. Liquid scintillation cocktail was added to this and radioactivity levels were measured with pellets corresponding to cell wall α-glucan fraction and supernatants to β-glucan-plus-galactomannan fraction. Third part of cell wall suspension was incubated with Quantazyme (MP Biomedicals; Q-Biogene) in 50mM potassium phosphate monobasic (pH 7.5), 60mM β-mercaptoethanol for 24 hr at 37°C. Radioactivity of pellets was measured after centrifugation and this corresponded to cell wall without β-1,3-glucan fraction, while supernatant was considered as β-1,3-glucan fraction. β-1,6-glucan was calculated as the remaining polysaccharide from total cell wall radioactivity minus radioactivity of galactomannan, α-glucan and β-1,3-glucan. All determinations were performed in duplicates, with at least three independent replicates for each strain (three independent experiments were performed for analysis at 25ºC and four independent experiments for the analysis at 34ºC).

### Immunoprecipitation and immunoblot analysis

Immunoprecipitation experiments were performed similar to that described in (Cortés et al., 2012). In brief, harvested cells were washed with stop solution (154mM NaCl, 10mM EDTA, 10mM NaN_3_, and 10mM NaF), followed by wash buffer (50mM Tris-HCl, pH 7.5, 5mM EDTA). Cells were then lysed by glass beads and eluted in lysis buffer (50mM Tris-HCl pH 7.5, 5mM EDTA, 200mM NaCl containing 100μM phenylmethylsulphonylfluoride, 1mM benzamidine and protease inhibitors - Complete EDTA-free, Roche Diagnostics). Cell lysate was clarified by centrifugation (4,500*g*, 1 min, 4°C). Cell membranes were separated from the supernatant thus obtained by centrifugation (16,000*g*, 1 hr, 4°C). The cell membrane pellet was resuspended in immunoprecipitation buffer (IPB; 50mM Tris-HCl, pH 7.5, 5mM EDTA, 200mM NaCl, 0.5% Tween 20, 100μM phenylmethylsulphonylfluoride, 1mM benzamidine and protease inhibitors - Complete EDTA-free, Roche Diagnostics), and agitated (1,300rpm, 30 min, 1°C; Thermomixer Comfort, Eppendorf). Solubilized membrane proteins were obtained in the supernatant from this agitated suspension by centrifugation (21,000*g*, 30 min, 4°C). The solubilized membrane protein supernatant fraction was further diluted with IPB, was incubated with antibody against GFP (Abcam) for 1 hr at 4°C. Sepharose protein A beads were added to this mix for 3 hr, later washed with IPB and boiled in sample buffer. Solubilized membrane proteins and IPs were then resolved on 4%-20% gels (Biorad), transferred to Immobilon-P membrane (Millipore), blocked and immunoblotted using monoclonal antibodies against GFP (1:2500, Abcam) or HA (1:5000, Roche Diagnostics). Peroxidase conjugated -rabbit or -mouse secondary antibodies (Jackson Laboratories) were used at 1:20000 dilutions. Signals were detected using enhanced chemiluminescence.

### Fluorescence distribution plotting

To quantify the distribution of Smi1p punctate structures along cell length, cells were imaged using *micro*LAMBDA spinning disk and maximum Z projections were used for analysis. In ImageJ, a line thick enough to cover the entire volume of the cell was drawn along the cell long axis and using the Plot Profile function of ImageJ plot values (V_1-n_) along the cell length values (L_1-n_) were copied on to Excel. The normalized cell length value was obtained using the formula l_1-n_, l_i_=L_i_/L_n_, with a value of 0 corresponding to one cell end and 1 to the other cell end. Similarly, normalized plot values along the cell length long axis were obtained as V_1-n_, V_i_=(V_i_/V_min_)/(V_max_-V_min_), with 0 being the lowest intensity value and 1 being the highest.

## 3. Results and discussion

### Multi-copy expression of *smi1*^*+*^ rescues *cps1-191* mutant defects

The Bgs1p temperature sensitive mutant, *cps1-191*, is defective in synthesis of the primary septum at the restrictive temperature of 36°C (Liu et al., 1999). At this temperature, mutants arrest as binucleate cells with a full unconstricted actomyosin ring. Failure in septum synthesis results in diminished support for the actomyosin ring and consequently, actomyosin rings are unstable, cannot constrict and are often observed to slide away from the cell middle (Arasada & Pollard, 2014; Cortés et al., 2015; Pardo & Nurse, 2003; Proctor et al., 2012). In brief, Bgs1p along with other glucan synthases are important both for septum synthesis and for providing adequate support to the cytokinetic machinery (Ramos et al., 2019).

To identify potential interacting partners for Bgs1p that may be involved in primary septum synthesis, we performed a multi-copy suppressor screen for *cps1-191* using a *S. pombe* genomic DNA plasmid library to identify DNA fragments that could reverse the colony formation defect of *cps1-191* mutant at the semi-permissive temperature 34°C (Sethi et al., 2016). We previously described Sbg1, a novel single-pass membrane protein whose over-expression suppressed *cps1-191* mutant phenotype (Sethi et al., 2016). This screen also identified *smi1*^*+*^ (SPBC30D10.17c) as a suppressor for *cps1-191* in four different instances (Figure 1A), which we characterize here. Multi-copy expression of Smi1p could not rescue *cps1-191* mutant at 36°C, establishing that Smi1p required a partially active Bgs1p/Cps1p to reverse the colony formation defect.

### Septum synthesis defect of *cps1-191* is partly corrected upon multi-copy expression of *smi1*^*+*^

The mutation in *cps1-191* affects the protein’s activity and localization required for septum synthesis (Cortés et al., 2015; Liu et al., 1999). We wanted to determine if overproduction of Smi1p in the mutant *cps1-191* background could suppress the septum synthesis defect of the mutant. Towards this goal, we shifted wild type cells with pEmpty (pAL), *cps1-191* cells carrying pEmpty and *cps1-191* cells carrying pSmi1 (pAL-*smi1*^+^) for 16hr at 34°C and stained them with aniline blue (marks the primary septum) and DAPI (marks the nuclei) (Figure 1B and 1C). Under these conditions, wild type cells maintained cylindrical morphology with binucleate cells displaying a medially placed normal division septum. However, only 17% of mutant *cps1-191* binucleate cells carrying pEmpty deposited a division septum. The cell morphology was also modified - the cells assumed a bulky and round morphology. In addition, 21% of the mutant cells accumulated multiple nuclei. However, upon overproduction of Smi1p in *cps1-191* background, the cell morphology was now improved and the cells were better able to synthesize a division septum. Up to 51% of binucleate cells had a clear division septum. In addition, upon overproduction of Smi1p, only 4% of the cells showed accumulation of multiple nuclei. This suggested that over production of Smi1p improved primary septum synthesis in the mutant *cps1-191* background.

### Overproduction of Smi1p in mutant *cps1-191* increases β-1,3-glucan level at the division septum

Bgs1p is responsible for the synthesis of the linear β-1,3-glucan at the division septum and this glucan forms part of the primary septum (Cortés et al., 2007). The absence or reduction of Bgs1 function results in an increase in α-glucan content in the cell wall as a compensatory mechanism (Cortés et al., 2007; Sethi et al., 2016). Also, the mutant cells deposit a much thicker secondary septum and the surrounding cell wall is also much thicker and less uniform as compared to wild type cells (Sethi et al., 2016). We considered if rescue of *cps1-191* by overproduction of Smi1p results in improved incorporation of linear β-1,3-glucan itself at the division site. A measure of the fluorescence intensity of calcofluor white (CW), a dye that specifically binds to linear β-1,3-glucan at the division septum, can be used to assess the levels of linear β-1,3-glucan (Cortés et al., 2007; Ribas & Cortés, 2016; Sethi et al., 2016). Towards this goal, cells from two different strains, one also labeled with FITC-concanavalin A, were analyzed together at 34°C (Figure 1D and 1E). FITC-concanavalin A binds to outer galactomannan layer of the cell wall and was used to differentiate between the two compared strains. Fluorescence intensity was calculated as arbitrary units using ImageJ and the final value was determined as intensity normalized to the length of the septum. Using fluorescence intensity of CW (normalized to septal length) in mutant *cps1-191* strain as a reference at 100%, we observed fluorescence intensity of wild type septa relative to *cps1-191* septa intensity to be 265%. Similarly, the CW intensity of septa in *cps1-191* cells overproducing Smi1p was 130% higher than the septum intensity of *cps1-191* mutant cells. Both these combinations of analyses were also performed in the reverse direction with the other strain being labeled with FITC-concanavalin A, and similar results for fluorescence intensity (fluorescence intensity of wild type septa relative to *cps1-191* septa intensity to be 265% and that of *cps1-191* cells overproducing Smi1p was 130% higher than the septum intensity of *cps1-191* mutant cells) were observed. Thus, overproduction of Smi1p indeed resulted in an increase in linear β-1,3-glucan levels at the division septum thus promoting better septum synthesis.

We had observed that upon overproduction of Smi1p, *cps1-191* cells showed higher levels of CW-stained material corresponding to linear β-1,3-glucan at the division septum. We thus speculated if multi-copy expression of *smi1*^*+*^ would also result in an overall improvement in ultrastructure of the cell wall in mutant *cps1-191* background as observed in the improvement of cell wall analysis and of septum linear β-1,3-glucan detected by CW staining. To this end, we imaged the mutant and the rescued cells using transmission electron microscopy. Similar to previous experiments, cells were shifted to 34°C for 16hr and then fixed for imaging (Figure 2). Wild type cells with pEmpty showed a clear three-layered division septum with an electron transparent primary septum flanked by electron dense secondary septum on either side. In addition, wild type cells showed a uniform cell wall. However, in the mutant *cps1-191* cells carrying pEmpty, a clear defect was observed in the cell wall ultrastructure. The cell wall appeared much thicker than wild type cells and also the division septum that was deposited did not display a clear electron transparent primary septum. A large proportion of the cells displayed a twisted or discontinuous primary septum as shown in the figure. However, upon multi-copy expression of *smi1*^*+*^, the surrounding cell wall was much better defined, and the thickness of the cell wall was comparable to that of the wild type cells. Also, the division septum was now restored to a three-layered structure, where a continuous primary septum could be clearly observed in most cells. In line with the observation of general improvement of septum and cell wall ultrastructure, cell wall fractionation analysis showed a partial restoration of the cell wall composition (Supplementary Data 1 and 2). Thus, it was evident that overproduction of Smi1p restored septum synthesis in the *cps1-191* mutant.

**Figure 2:**
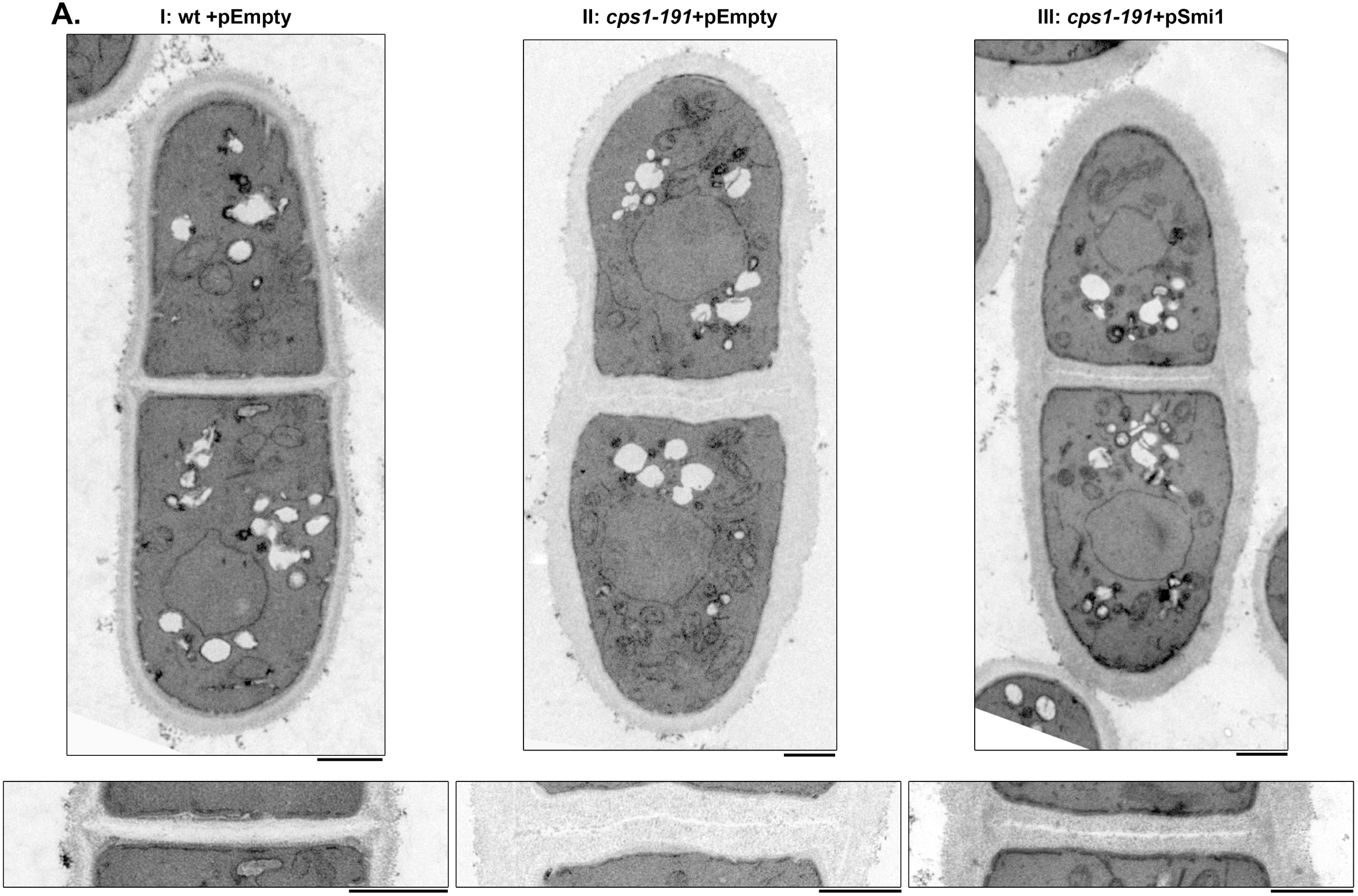
Multi-copy expression of *smi1*^*+*^ improves cell wall defects of the glucan synthase *cps1-191* mutant. (A) TEM images of septated cells after 16 hr at 34°C from the following strains, I: wt+pEmpty (MBY8558), II: *cps1-191+*pEmpty (MBY8944) and III: *cps1-191+*pSmi1 (MBY8945). Images at the bottom display the same division septum at the same a higher magnification. Scale bar 1μm.

**Figure 3:**
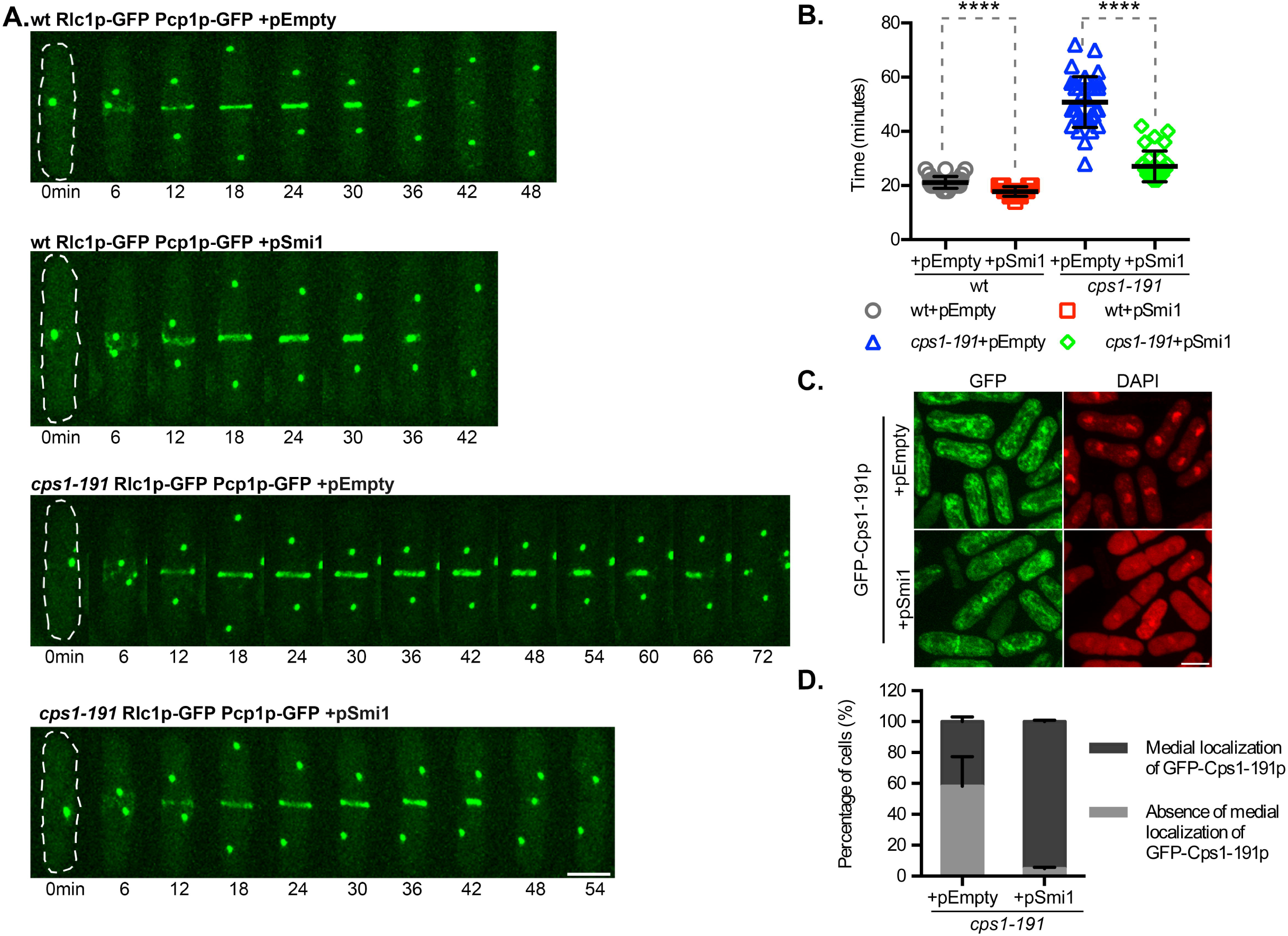
Multi-copy expression of *smi1*^*+*^ improves *cps1-191* mutant actomyosin ring and division defects. (A) Time-lapse maximum Z projection spinning disk confocal montages of actomyosin ring in the indicated strains: wt Rlc1p-GFP Pcp1p-GFP+pEmpty (MBY9493), wt Rlc1p-GFP Pcp1p-GFP+pSmi1 (MBY9510), *cps1-191* Rlc1p-GFP Pcp1p-GFP+pEmpty (MBY9454) and *cps1-191* Rlc1p-GFP Pcp1p-GFP+pSmi1 (MBY9506) after 3.5 hr at 34°C. 0min indicates time of spindle pole body duplication. Green, Rlc1p-GFP Pcp1p-GFP. (B) Graph shows the time taken (in minutes) for the ring to constrict in the indicated strains in (A). p-value<0.0001 indicated by **** (two-tailed T-test, n≥30cells). (C) Maximum Z projection spinning disk confocal images of indicated strains: GFP-Cps1-191p+pEmpty_his3 (MBY9188) and GFP-Cps1-191p+pSmi1_his3 (MBY9199) after 6 hr at 34°C. Cells were fixed and stained with DAPI to identify binucleate cells. (D) Quantification for the presence or absence of medial GFP-Cps1-191p localization at 34°C for experiment in (C). (n=3, ≥100cells). Scale bar 5μm.

### Improved actomyosin ring dynamics in *cps1-191* mutant upon overproduction of Smi1p

*cps1-191* is unable to constrict its actomyosin ring at the restrictive temperature of 36°C, possibly as a result of lack of support from an ingressing division septum (Liu et al., 1999). We considered if at the lower restrictive temperature of 34°C, *cps1-191* would present a similar actomyosin ring constriction defect. To this end, we imaged *cps1-191* cells expressing Rlc1p-GFP (myosin II regulatory light chain protein to mark the actomyosin ring) and Pcp1p-GFP (spindle pole body marker to judge mitotic progression) harboring the empty plasmid and imaged actomyosin ring dynamics (Figure 3A and 3B). We observed that almost all the cells were able to constrict their actomyosin rings (average constriction time 50.83 ± 9.37 min), though much slower than wild type cells (average constriction time 21.12 ± 2.17 min) imaged under the same conditions. We determined the ring constriction time in wild type cells with pSmi1 to be an average of 17.86 ± 1.73 min. We imaged *cps1-191* cells expressing Rlc1p-GFP Pcp1p-GFP with pSmi1 in a similar fashion to determine if multi-copy expression of *smi1*^*+*^ had an impact on ring dynamics of *cps1-191* mutant. Interestingly, we observed that indeed the ring dynamics were now improved upon overproduction of Smi1p (average constriction time 32.67 ± 6.39 min). In conclusion, overproduction of Smi1p in mutant *cps1-191* cells improved both ring dynamics and septum synthesis efficiency. It is possible that the actomyosin ring dynamics in the mutant cells were improved as a consequence of some contributions from an ingressing division septum upon multi-copy expression of *smi1*^*+*^.

### Overproduction of Smi1p improves retention of mutant protein Cps1-191p to the division site

Previous work has shown that the product of the *cps1-191* mutant does not localize correctly to the division site, with the bulk of it being retained at the endoplasmic reticulum (ER), even at the permissive temperature of 24°C (Cortés et al., 2015; Sethi et al., 2016). We addressed if overproduction of Smi1p in *cps1-191* mutant rescued the localization defect of the mutant protein Cps1-191p and this in turn could possibly explain the improved ring dynamics and division septum assembly. We thus compared *GFP-cps1-191* cells (which express Cps1-191p fused to GFP) carrying either pEmpty or pSmi1, grown at 34°C (Figure 3C and 3D). Cells were fixed and stained with DAPI to identify binucleate cells, as at this stage Bgs1p can be detected at the division site in wild type cells. GFP-Cps1-191p strain expressing pEmpty failed to show medial localization of the mutant Cps1-191p in ∼58.24% ± 19.16% of binucleate cells and was mostly retained at the ER. However, upon multi-copy expression of *smi1*^*+*^, the mutant protein localization to the division site was improved and almost all the binucleate cells (95.16% ± 0.91%) showed medial localization of GFP-Cps1-191p in this background. Thus, overproduction of Smi1p promoted the correct localization of the mutant protein to the division site, thereby potentially reversing the primary septum assembly defect.

### Smi1p is essential for cell viability

The *S. pombe* database suggests that Smi1p is an essential gene (Hayles et al., 2013; Kim et al., 2010; Wood et al., 2012). To ascertain this, we obtained heterozygous diploids with one wild type copy of *smi1*^*+*^ and the other copy deleted with a KanMX cassette (causing G418 resistance). On tetrad dissection analysis, we observed a 2:2 growth pattern (Figure 4A). All the viable colonies were confirmed to be wild type cells and G418-sensitive. Thus, we conclude that *smi1*^*+*^ is essential for cell viability. We next analysed *smi1*Δ spores germinated from this diploid. When the *smi1Δ* spores from this diploid were cultured, they failed to germinate even after 30 hrs of incubation in rich medium. Thus, Smi1p is essential for spore germination.

**Figure 4:**
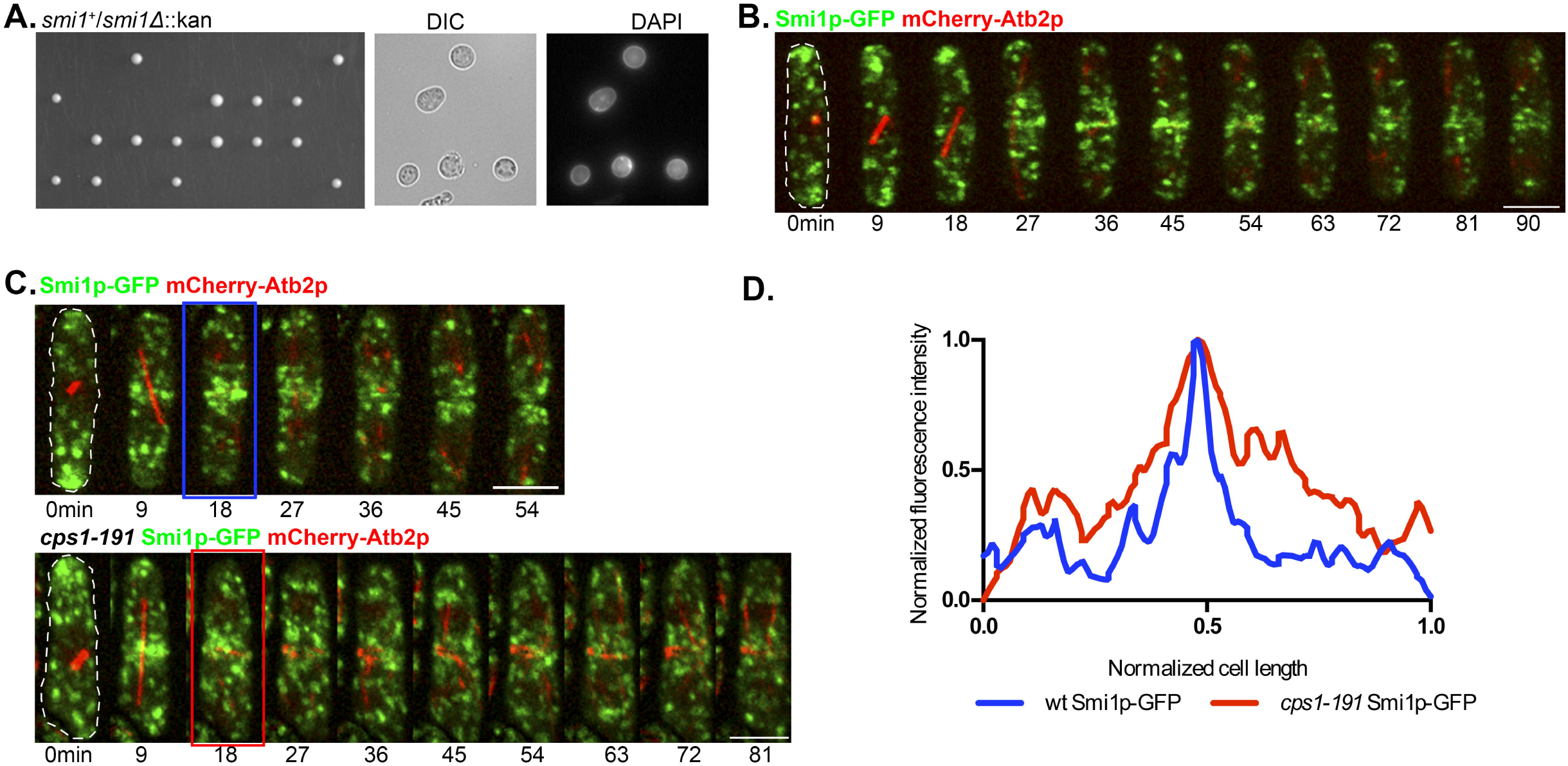
Characterization of Smi1p. (A) Tetrad dissection analysis of diploid *smi1*Δ*/smi1*^*+*^ (MBY9158) showing 2:2 segregation of growth on YES plates. Images show germinated *smi1*Δ spores. (B) Time-lapse maximum Z projection spinning disk confocal montage of the indicated strain Smi1p-GFP mCherry-Atb2p (MBY8651). Green, Smi1p*-*GFP. Red, mCherry-Atb2p. 0min marks spindle pole body duplication as judged by a short microtubule spindle. (C) Time-lapse maximum Z projection spinning disk confocal montage of the indicated strains: Smi1p*-*GFP mCherry-Atb2p (MBY8651) and *cps1-191* Smi1p*-*GFP mCherry-Atb2p (MBY9133) after 2 hr at 36°C. (D) Plot compares fluorescence intensity of Smi1p along cell length between wild-type cells and *cps1-191* cells at spindle breakdown at 36°C (indicated by blue and red boxed in the montages in (C)). Scale bar 5μm.

### Smi1p localizes to punctate structures

To further understand the role played by Smi1p in cytokinesis, we generated a strain in which Smi1p was tagged with GFP at the C-terminus. We observed Smi1p-GFP to localize to punctate structures throughout the cell (Figure 4B). We analysed the localization of Smi1p through the cell cycle by using mCherry-Atb2p (tubulin) as a cell cycle marker. Smi1p punctae were enriched at the cell ends with some structures also present in the cytoplasm. In cells with a long microtubule spindle, we observed that Smi1p punctae were largely concentrated at the division site. The role of Smi1p at the division site is unknown, however the localization is consistent with that of *S. cerevisiae* Knr4p/Smi1p at the bud neck at cytokinesis. It is possible that Smi1p associates with other cytokinetic proteins or cell wall remodelling proteins during cytokinesis.

It was evident from localization of Smi1p in wild type cells that Smi1p was enriched both at the cell ends and at the division site. We speculated if the enrichment of Smi1p to the division site was dependent on functional Bgs1p. To this end, we imaged *cps1-191* Smi1p-GFP mCherry-Atb2p at the restrictive temperature of 36°C after shift up for 2 hr (Figure 4C and 4D). The Smi1p punctae appeared smaller and more diffused in the cytoplasm, but most importantly we compared the distribution of fluorescence intensity of Smi1p along the cell length after spindle disassembly both in wild type and mutant cells at 36°C. In wild type cells, we observed a sharp peak of fluorescence intensity of Smi1p corresponding to highly concentrated Smi1p puncta at the division site. However, in the mutant *cps1-191* background, though Smi1p puncta were enriched at the cell middle, the fluorescence intensity was more spread out in 58%±12% of the cells as compared to that in wild type cells. It is possible that proper localization of Smi1p at the division site requires the activity of functional and correctly localized Bgs1p or any of the proteins that interact with Bgs1p at the division site.

### Bgs1p and Smi1p localization is associated

As *smi1*^*+*^ was identified as a genetic suppressor of the *cps1-191* mutant, we considered that the two proteins may localize at the same cellular compartments to function cooperatively. As such, we imaged a strain co-expressing GFP-Bgs1p and Smi1p-mCherry. As shown in Figure 5A, Smi1p and Bgs1p localized to the cell ends and to the division site at similar time points over the cell cycle. The colocalization was more prominent in the cytoplasm at interphase where Bgs1p clearly was present in many of the Smi1p punctate structures. At mitosis, although both Bgs1p and Smi1p localized to the cell middle and septum membrane, the localization was not as strongly correlated. It is possible that Smi1p and Bgs1p are transported together to the division site but upon arriving at the division site only Bgs1p is retained for its activity in septum synthesis.

**Figure 5:**
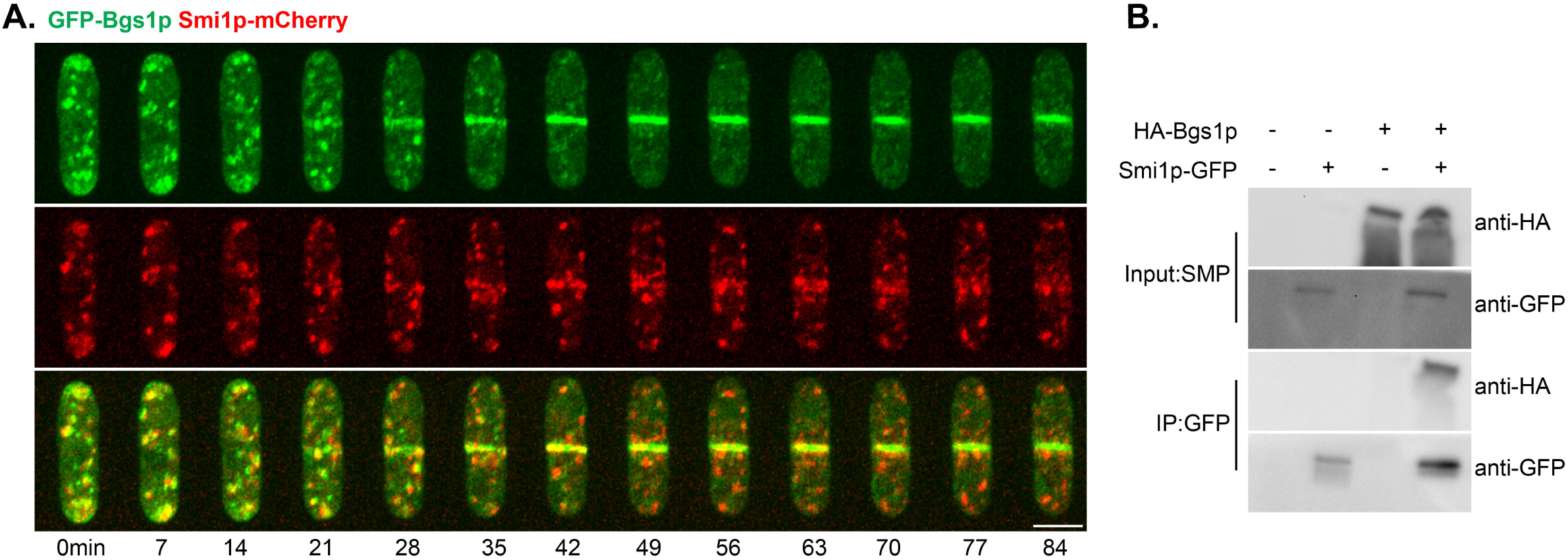
Smi1p and Bgs1p colocalize to cell ends and septum and are associated at the plasma membrane. (A) Time-lapse maximum Z projection spinning disk confocal montage of the indicated strain GFP-Bgs1p Smi1p-mCherry (MBY9184). Scale bar 5μm. (B) Physical interaction between Bgs1p and Smi1p. Solubilized membrane proteins from the indicated strains: wt (MBY192), Smi1p*-*GFP (MBY8650), HA-Bgs1p (MBY8702) and HA-Bgs1p Smi1p*-*GFP (MBY8709) were immunoprecipitated (IP) with anti-GFP antibodies. Solubilized membrane proteins (input, top) and IP (bottom) were transferred to the same membrane and blotted with monoclonal anti-HA antibodies and monoclonal anti-GFP antibodies.

### Smi1p and Bgs1p physically interact

Given our findings, we speculated if overproduction of Smi1p improved the defects of the mutant by direct interaction with the Bgs1p protein. To this end, we assessed if Smi1p and Bgs1p physically interacted with each other in wild type cells. We used the strain co-expressing Smi1p-GFP and HA-Bgs1p for this experiment. Upon immunoprecipitating Smi1p-GFP using anti-GFP antibodies, we were able to detect HA-Bgs1p in the pull-down complex (Figure 5B). However, a band corresponding to HA-Bgs1p was not detected in any of the control strains. Thus, we conclude that either Smi1p physically interacts with Bgs1p or that both localize to the same plasma membrane domains.

## 4. Discussion

In conclusion, in this study we have isolated a suppressor, Smi1p, for the septum synthesis defective temperature sensitive mutant *cps1-191* and provide an initial characterization of this protein. This previously uncharacterized essential protein interacts with Bgs1p and upon overproduction in the mutant *cps1-191* background promotes retention of the mutant Cps1-191p protein at the division site, improves ring dynamics and cell wall ultrastructure. The Smi1p protein localizes to punctate structures in the cytoplasm and its localization at the division site is dependent on functional Bgs1p. We hypothesize that Smi1p and Bgs1p are transported to the cell division site together or Smi1p contributes to the transport of Bgs1p to the cell division site. In the future, identification of potential interacting partners of Smi1p or detailed analysis of phenotype of a Smi1p temperature-sensitive mutant, would provide more insight into the molecular mechanism by which overproduction of Smi1p could rescue *cps1-191*.

In a previous work, we have shown that *cps1-191* is rescued upon overproduction of the single-pass transmembrane protein Sbg1p. Given the ability of these two proteins to rescue the *cps1-191* allele (D277N), upon overproduction, it is possible that Bgs1p, Sbg1p, and Smi1p are present in a single complex which facilitates Bgs1p transport to and/or retention and stabilization at the division site. Alternatively, it is possible that *cps1-191* is defective in interaction with an as yet unidentified protein involved in its retention at the division site, but that it can also be retained at the division site through other weak interactors that bind the protein elsewhere (i.e. not in the vicinity of D277) and Sbg1p and Smi1p might collaborate with these weak interactors. Identification of the exact binding sites of Sbg1p and Smi1p on Bgs1p will help distinguish between these models. Since D277 is in the cytoplasmic domain 1 out of 8 of Bgs1 (http://wlab.ethz.ch/protter/#up=BGS1_SCHPO&tm=auto) and this residue forms salt bridges with R1151 on cytoplasmic domain 4 out of 8 of Bgs1p containing its UDP-glucose binding domain, it is also possible that Sbg1 and Smi1 overproduction stimulate or stabilize the residual activity of Bgs1-191p (product of *cps1*-*191*) (https://alphafold.ebi.ac.uk/entry/Q10287; Jumper et al., 2021; Liu et al., 1999).

Recently Wu and colleagues identified Smi1p in a proteomic screen for Bgs4p physical interactors (Longo et al., 2022). Interestingly, although they found synthetic lethality between temperature-sensitive *cps1-191* and *cwg1-2* (a Bgs4p mutant), in mistargeting experiments it was noticed that Smi1p effect was stronger on Bgs4p and Ags1p probably because of their essential function in cell integrity. A combination of our work and that of Wu and colleagues suggests that Smi1p is involved in the localization and/or activation of more than one β-1,3-glucan synthase. The molecular mechanisms of β-1,3-glucan transport and activation via Sbg1p and Smi1p can be investigated in the coming years.

## Acknowledgements

We thank Dhivya Subramanian, Huang Yinyi, Evelyn Tao, Rajesh Patkar and Zhang Dan for their useful suggestions and ideas. We also thank current and past MB and NN lab members for their guidance.

## Author Contributions

KS, JCGC and MS performed the experiments. KS and MB prepared the manuscript. MO, NN, JCR and MB supervised the study.

## Funding

MB was funded by core funds from Temasek Life Sciences Laboratory, Wellcome Trust SIA (WT101885MA). NN was funded by core funds from Temasek Life Sciences Laboratory. JCR was supported by research funds from the Spanish Ministry of Science and Innovation (MICINN, Spain) and the European Regional Developmental Fund (FEDER, EU) (PGC2018-098924-B-I00), and the Regional Government of Castile and Leon (JCYL, Spain) and the European Regional Developmental Fund (FEDER, EU) (CSI150P20 and “Escalera de Excelencia” CLU-2017-03), and MO was supported by research funds from Bio-imaging Center (S0991205) of Japan Women’s University established as a private university in Japan with the support of Ministry of Education, Culture, Sports Science and Technology.

## Conflict of Interest

The authors declare that the research was conducted in the absence of any commercial or financial relationships that could be construed as a potential conflict of interest.

## Notes

### Competing Interest Statement

The authors have declared no competing interest.

