## Supplemental Material for "Fission yeast Smi1p participates in the synthesis of the primary septum by regulating β-1,3-glucan synthase Bgs1p function"

### *Supplementary Material*

**Supplementary Data 1: Incorporation at 25ºC of [$^{14}$C]glucose into cell wall polysaccharides.**

**Incorporation at 25ºC of [$^{14}$C]glucose into cell wall polysaccharides. % in the cell**

% Incorporation of [$^{14}$C]glucose in the cell[b]

| Strain[a] | Cell wall[c] | α-glucan[c] | β(1,3)-glucan[c] | β(1,6)-glucan[c] | Galactomannan[c] |
|---|---|---|---|---|---|
| Wild type pAL<br>MBY8558 | 30.3 ± 1.6 | 8.2 ± 0.8 | 16.6 ± 1.0 | 1.3 ± 0.4 | 4.2 ± 0.7 |
| *cps1-191* pAL<br>MBY8944 | 36.9 ± 1.1 | 12.0 ± 0.5 | 17.7 ± 0.7 | 2.2 ± 0.6 | 5.0 ± 0.8 |
| *cps1-191* pAL-*smi1*$^{+}$<br>MBY8945 | 34.9 ± 1.5 | 10.6 ± 0.4 | 17.6 ± 0.7 | 2.4 ± 0.3 | 4.3 ± 0.3 |

[a] Early log-phase cell cultures were grown in MM at 25ºC. [$^{14}$C]glucose was added 24 hours before harvesting.

[b] Percentage incorporation of [$^{14}$C]glucose = cpm incorporated per fraction x 100/total cpm incorporated into the cell. Values are the means and SDs calculated from three independent experiments.

[c] Values are percentages of the cell wall and the corresponding polysaccharide in the cell.

**Incorporation at 25ºC of [$^{14}$C]glucose into cell wall polysaccharides. % in the cell wall**

% Incorporation of [$^{14}$C]glucose in the cell wall[b]

| Strain[a] | Cell wall[c] | α-glucan[c] | β(1,3)-glucan[c] | β(1,6)-glucan[c] | Galactomannan[c] |
|---|---|---|---|---|---|
| Wild type pAL MBY8558 | 100 | 27.1 ± 1.9 | 54.7 ± 1.0 | 4.3 ± 1.4 | 13.9 ± 1.7 |
| *cps1-191* pAL MBY8944 | 100 | 32.4 ± 0.4 | 47.8 ± 1.2 | 6.1 ± 1.2 | 13.7 ± 2.1 |
| *cps1-191* pAL-*smi1*$^{+}$ MBY8945 | 100 | 30.4 ± 0.4 | 50.5 ± 1.2 | 6.8 ± 1.3 | 12.3 ± 0.9 |

[a] Early log-phase cell cultures were grown in MM at 25º. [$^{14}$C]glucose was added 24 hours before harvesting.

[b] Percentage incorporation of [$^{14}$C]glucose = cpm incorporated per fraction x 100/total cpm incorporated into the cell wall. Values are the means and SDs calculated from three independent experiments.

[c] Values are percentages of the corresponding polysaccharide in the cell wall.

**Supplementary Data 2: Incorporation at 34ºC of [$^{14}$C]glucose into cell wall polysaccharides.**

**Incorporation at 34ºC of [$^{14}$C]glucose into cell wall polysaccharides. % in the cell**

| | % Incorporation of [$^{14}$C]glucose in the cell[b] | | | | |
|---|---|---|---|---|---|
| Strain[a] | Cell wall[c] | α-glucan[c] | β(1,3)-glucan[c] | β(1,6)-glucan[c] | Galactomannan[c] |
| Wild type pAL MBY8558 | 26.2 ± 2.4 | 6.5 ± 0.9 | 15.4 ± 0.9 | 1.1 ± 0.4 | 3.2 ± 0.4 |
| *cps1-191* pAL MBY8944 | 37.5 ± 3.4 | 14.4 ± 1.5 | 17.9 ± 1.5 | 2.9 ± 0.5 | 2.3 ± 0.4 |
| *cps1-191* pAL-*smi1*$^{+}$ MBY8945 | 35.2 ± 3.0 | 12.7 ± 1.2 | 18.5 ± 1.6 | 1.7 ± 0.5 | 2.3 ± 0.3 |

[a] Early log-phase cell cultures were grown in MM at 25ºC and shifted to 34ºC for 16 hours. [$^{14}$C]glucose was added 16 hours before harvesting.

[b] Percentage incorporation of [$^{14}$C]glucose = cpm incorporated per fraction x 100/total cpm incorporated into the cell. Values are the means and SDs calculated from four independent experiments.

[c] Values are percentages of the cell wall and the corresponding polysaccharide in the cell.

**Incorporation at 34ºC of [$^{14}$C]glucose into cell wall polysaccharides. % in the cell wall**

% Incorporation of [$^{14}$C]glucose in the cell wall[b]

| Strain[a] | Cell wall[c] | $\alpha$-glucan[c] | $\beta$(1,3)-glucan[c] | $\beta$(1,6)-glucan[c] | Galactomannan[c] |
|---|---|---|---|---|---|
| Wild type pAL MBY8558 | 100 | 24.9 $\pm$ 1.3 | 58.8 $\pm$ 1.8 | 4.1 $\pm$ 1.2 | 12.2 $\pm$ 0.6 |
| *cps1-191* pAL MBY8944 | 100 | 38.4 $\pm$ 1.7 | 47.7 $\pm$ 1.3 | 7.7 $\pm$ 1.4 | 6.2 $\pm$ 0.9 |
| *cps1-191* pAL-*smi1*$^+$ MBY8945 | 100 | 36.1 $\pm$ 2.0 | 52.4 $\pm$ 1.8 | 4.9 $\pm$ 0.6 | 6.6 $\pm$ 0.3 |

[a] Early log-phase cell cultures were grown in MM at 25ºC and shifted to 34ºC for 16 hours. [$^{14}$C]glucose was added 16 hours before harvesting.

[b] Percentage incorporation of [$^{14}$C]glucose = cpm incorporated per fraction x 100/total cpm incorporated into the cell wall. Values are the means and SDs calculated from four independent experiments.

[c] Values are percentages of the corresponding polysaccharide in the cell wall.

**Table S1** ***S. pombe*** **strains**

| | | |
|---|---|---|
| MBY 102 | *ade6-210 ura4-Δ18 leu1-32 $h^+$* | Lab collection |
| MBY 103 | *ade6-216 ura4-Δ18 leu1-32 $h^-$* | Lab collection |
| MBY 192 | *ura4-Δ18 leu1-32 $h^-$* | Lab collection |
| MBY 1148 | *cps1-191 ade6-M21x ura4-Δ18 leu1-32 $h^+$* | Lab collection |
| MBY 5730 | *cps1-191 $rlc1^+$-gfp::$ura^+$ $pcp1^+$-gfp:KanMX6* | Lab collection |
| MBY 5732 | *$rlc1^+$-gfp::$ura^+$ $pcp1^+$-gfp:KanMX6 $h^-$* | Lab collection |
| MBY 8558 | *ura4-Δ18 leu1-32 with pEmpty $h^-$* | Lab collection |
| MBY 8650 | *smi1-GFP:kanMX6 leu1-32 ura4-Δ18 $h^-$* | This study |
| MBY 8651 | *smi1-GFP:kanMX6 mCherry-$atb2^+$:hph leu1-32 ura4-Δ18* | This study |
| MBY 8702 | *$P_{bgs1+}$::3XHA-$bgs1^+$:$leu1^+$ bgs1Δ::$ura4^+$ leu1-32 ura4-Δ18 $h^+$* | Lab collection |
| MBY 8709 | *$smi1^+$-GFP:kanMX6 $P_{bgs1+}$::3XHA-$bgs1^+$:$leu1^+$ bgs1Δ::$ura4^+$ leu1-32 ura4-Δ18 $h^-$* | This study |
| MBY 8944 | *cps1-191 ade6-M21x ura4-Δ18 leu1-32 with pEmpty $h^+$* | This study |
| MBY 8945 | *cps1-191 ade6-M21x ura4-Δ18 leu1-32 with pSmi1 $h^+$* | This study |
| MBY 8947 | *cps1-191 ade6-M21x ura4-Δ18 leu1-32 with pCps1 $h^+$* | This study |

| | | |
|---|---|---|
| MBY 9100 | *$h^+/h^+$ smi1Δ:kanMX/$sbg1^+$ade6-M210/ade6-M216 ura4-Δ18/ura4-Δ18 leu1-32/leu1-32* | Bioneer Korea |
| MBY 9133 | *cps1-191 $smi1^+$-GFP:kanMX6 mCherry-$atb2^+$:hph ura4-Δ18 leu1-32 ade6-M21x* | This study |
| MBY 9158 | *$h^+/h^-$ smi1Δ:kanMX/$sbg1^+$ ade6-M210/ade6-M216 ura4-Δ18/ura4-Δ18 leu1-32/leu1-32* | This study |
| MBY 9184 | *$smi1^+$-mCherry-natMX6 $P_{bgs1+}$::GFP-$bgs1^+$: $leu1^+$ bgs1Δ::$ura4^+$ ade6-210 ura4-D18 leu1-32 $h^-$* | This study |
| MBY 9188 | *$P_{bgs1+}$::GFP-12A-cps1-191: $leu1^+$ bgs1Δ::$ura4^+$ leu1-32 ura4-Δ18 his3-Δ1 with pEmpty_his3 $h^-$* | This study |
| MBY 9199 | *$P_{bgs1+}$::GFP-12A-cps1-191: $leu1^+$ bgs1Δ::$ura4^+$ leu1-32 ura4-Δ18 his3-Δ1 with pSmi1_his3 $h^-$* | This study |
| MBY 9454 | *cps1-191 $rlc1^+$-gfp::$ura^+$ $pcp1^+$-gfp:KanMX6 with pEmpty $h^+$* | This study |
| MBY 9493 | *$rlc1^+$-gfp::$ura^+$ $pcp1^+$-gfp:KanMX6 with pEmpty $h^-$* | This study |
| MBY 9506 | *cps1-191 $rlc1^+$-gfp::$ura^+$ $pcp1^+$-gfp:KanMX6 with pSmi1 $h^+$* | This study |
| MBY 9510 | *$rlc1^+$-gfp::$ura^+$ $pcp1^+$-gfp:KanMX6 with pSmi1 $h^+$* | This study |

**Table S2 Plasmids used in this study**

| | |
|---|---|
| pCDL1522 | p*Hyg-eGFP:$Amp^r$* |
| pCDL1000 | p*AL-KS-empty* (referred to as pEmpty) |

| | |
|---|---|
| pBac15 | p*AL-KS-bgs1*⁺ (referred to as pBgs1) |
| pBac16 | p*AL-KS-smi1*⁺ (referred to as pSmi1) |
| pBac6 | p*AL-KS-empty:his3*⁺ (referred to as pEmpty_his3) |
| pBac8 | p*AL-KS-smi1⁺:his3*⁺ (referred to as pSmi1_his3) |

**Table S3 Primers used in this study**

| | |
|---|---|
| MOH 1207 | AATTAACCCTCACTAAAGGG |
| MOH 1208 | GTAATACGACTCACTATAGGGC |
| MOH 5904 | GTGTCCCAAAACAGCACTATTGCCGAAGCTAGTTCTCTTCAAGCTCAAGAAGAGGAAAAAGAAATAGAGACTACAAGTGTGAAGCAAGAGATCCCCGGGTTAATTAACAG |
| MOH 5905 | TTTACAAGATAGAGATAGACAAGTGCACTAGGAACAAGAAGGCACATAATATATCTTCTGAGGCGGTCATCATCACACTGAATGTCTAGCGCGGCCGCATAGGCCACTAG |
